# WormQTL2: an interactive platform for systems genetics in *Caenorhabditis elegans*

**DOI:** 10.1101/766386

**Authors:** Basten L. Snoek, Mark G. Sterken, Margi Hartanto, Albert-Jan van Zuilichem, Jan E. Kammenga, Dick de Ridder, Harm Nijveen

## Abstract

Quantitative genetics provides the tools for linking polymorphic loci (QTLs) to trait variation. Linkage analysis of gene expression is an established and widely applied method, leading to the identification of expression quantitative trait loci (eQTLs). (e)QTL detection facilitates the identification and understanding of the underlying molecular components and pathways, yet (e)QTL data access and mining often is a bottleneck. Here we present WormQTL2 (www.bioinformatics.nl/WormQTL2/), a database and platform for comparative investigations and meta-analyses of published (e)QTL datasets in the model nematode worm *C. elegans*. WormQTL2 integrates six eQTL studies spanning 11 conditions as-well-as over 1000 traits from 32 studies and allows experimental results to be compared, reused, and extended upon to guide further experiments and conduct systems-genetic analyses. For example, one can easily screen a locus for specific cis-eQTLs that could be linked to variation in other traits, detect gene-by-environment interactions by comparing eQTLs under different conditions, or find correlations between QTL profiles of classical traits and gene expression.

## Introduction

The nematode *Caenorhabditis elegans* has been instrumental as a model organism in studying genotype-phenotype relationships. Its genetic tractability in combination with a rapid life cycle and a large body of experimental data provides a powerful platform for investigating the genetics of complex traits. Extensive molecular, cellular, and physiological insights have been obtained using knockout mutants, RNAi-treatment, and various other techniques carried out in the canonical genotype Bristol N2 (1). Yet, due to many selection bottlenecks in the laboratory, the Bristol N2 strain has become the “lab worm” and is not representative of the effect of wild type alleles, as reviewed by (1). Over the last decade, the study of natural variation in *C. elegans* has made rapid progress, leading to the identification of over a dozen allelic variants contributing to natural phenotypic variation (2–26). Most quantitative genetics studies in this model animal have been conducted in recombinant inbred line (RIL) panels derived from two polymorphic wild type strains, namely N2 (isolated in Bristol, UK) and CB4856 (isolated in Hawaii, USA) (1,27–29). A substantial amount of phenotypic, genotypic, and high-throughput molecular data has been gathered across these recombinant inbred panels, as well as from introgression lines (30, 31) (31: PREPRINT NOT PEER REVIEWED), and many other wild isolates (32–34); for a more detailed overview, see reviews (1, 35). Together, the field of quantitative genetics in *C. elegans* has been very productive. However, accessing all this data, and using it for follow-up studies or for comparative analysis can be challenging. Furthermore, recent in-depth studies on the ecology of *C. elegans* yielded even more phenotypic and genotypic information on novel wild isolates (34,36–46). Inclusion of diverse genetic backgrounds in experiments is therefore a welcome and useful addition to the work on the canonical N2 strain as it can identify novel modifiers of well-studied pathways and thereby shed light on molecular mechanisms of genetic variation (46–52). A collection of data on wild isolates, including genome sequences, is curated and available via the *C. elegans* Natural Diversity Resource (CeNDR) (33).

The genetics of complex traits can be unravelled by performing Quantitative Trait Locus (QTL) analysis. QTLs are parts of the genome that harbour genetic variation associated with trait variation measured between different genotypes. Most QTL studies in *C. elegans* make use of recombinant inbred lines (RILs) derived from Bristol N2 and the genetically divergent Hawaiian strain CB4856 (28, 29). Traits such as body size, fecundity, aging, or pathogen sensitivity have been linked to underlying loci by QTL analysis (11,53–59). Moreover, gene expression studies comparing N2 and CB4856 show ample genotype dependent gene expression variation (60, 61). When microarray platforms became affordable and more easily usable, as currently has happened for RNA-seq, the range of phenotypes in RIL populations was extended with genome-wide gene expression. The identified expression QTLs (eQTLs) are - like classical QTLs - polymorphic loci linked to gene expression variation (62, 63). eQTLs can be *cis* or *trans*-acting: a *cis-*eQTL is normally defined as an eQTL mapping to the genomic location of the gene which it affects (usually within 1-2 Mb for *C. elegans*), while for a *trans*-eQTL genetic variation in another genomic region causes the change in gene expression. Both *cis*- and *trans-*eQTLs can shed light on the regulatory mechanisms underlying variation in molecular and phenotypic traits. Furthermore, the co-localization of *trans*-eQTLs, coined *trans*-bands or eQTL hotspots, makes them interesting for regulatory network analysis, as it is assumed that one or a few polymorphic ‘master regulator’ genes affect the expression of many target genes. In principle, by connecting each gene to its regulators, gene regulatory networks can be constructed from these eQTLs (29,64–71).

The identification of causal genes underlying *trans*-bands in *C. elegans* can be challenging. One of the complicating aspects is that the ultimate causal variant may act indirectly (*e.g.* through behaviour or hormone), rather than via a direct route (*e.g.* a transcription factor) (3). To date, only two such causal variants have been experimentally confirmed: *npr-1* and *amx-2* (3, 50). Another limitation is the still relatively low resolution of current eQTL analysis, typically yielding eQTLs spanning large genomic regions with hundreds of genes. Therefore, combining data from different experiments will result in better contextual information leading to a more detailed reconstruction of the regulatory mechanisms and more specific candidate gene lists.

Here, we present WormQTL2 (www.bioinformatics.nl/WormQTL2/), the platform for systems genetics in *C. elegans*. WormQTL2 is based on a versatile and interactive analysis platform for *Arabidopsis* QTL data called AraQTL, www.bioinformatics.nl/AraQTL (72). We used this framework combined with the ideas from WormQTL (www.WormQTL.org), (73–75) to present the *C. elegans* QTL data in an interactive manner. The WormQTL2 interface improves on WormQTL (73–75), allowing for dynamic and interactive cross-study analyses to aid hypothesis driven genotype-phenotype investigations. Through WormQTL2, data on natural variation in *C. elegans* and the identified QTLs have been made accessible and interactively approachable, beyond the original publications. The majority of the data in WormQTL2 are expression QTLs based on data from six different *C. elegans* eQTL studies (Table 1) (28,50,76–79). Moreover, phenotypic QTLs are well represented with data on more than 1000 traits from 32 studies. This extends the exploratory options by integrating QTL data on classical phenotypes, gene expression levels as well as protein- and metabolite levels (2,3,13,23,30,34,41,47,49,53–59,61,76,80–90) (Table 2). In this paper, we present WormQTL2 and showcase its use by presenting short research scenarios.

**Table 1:**
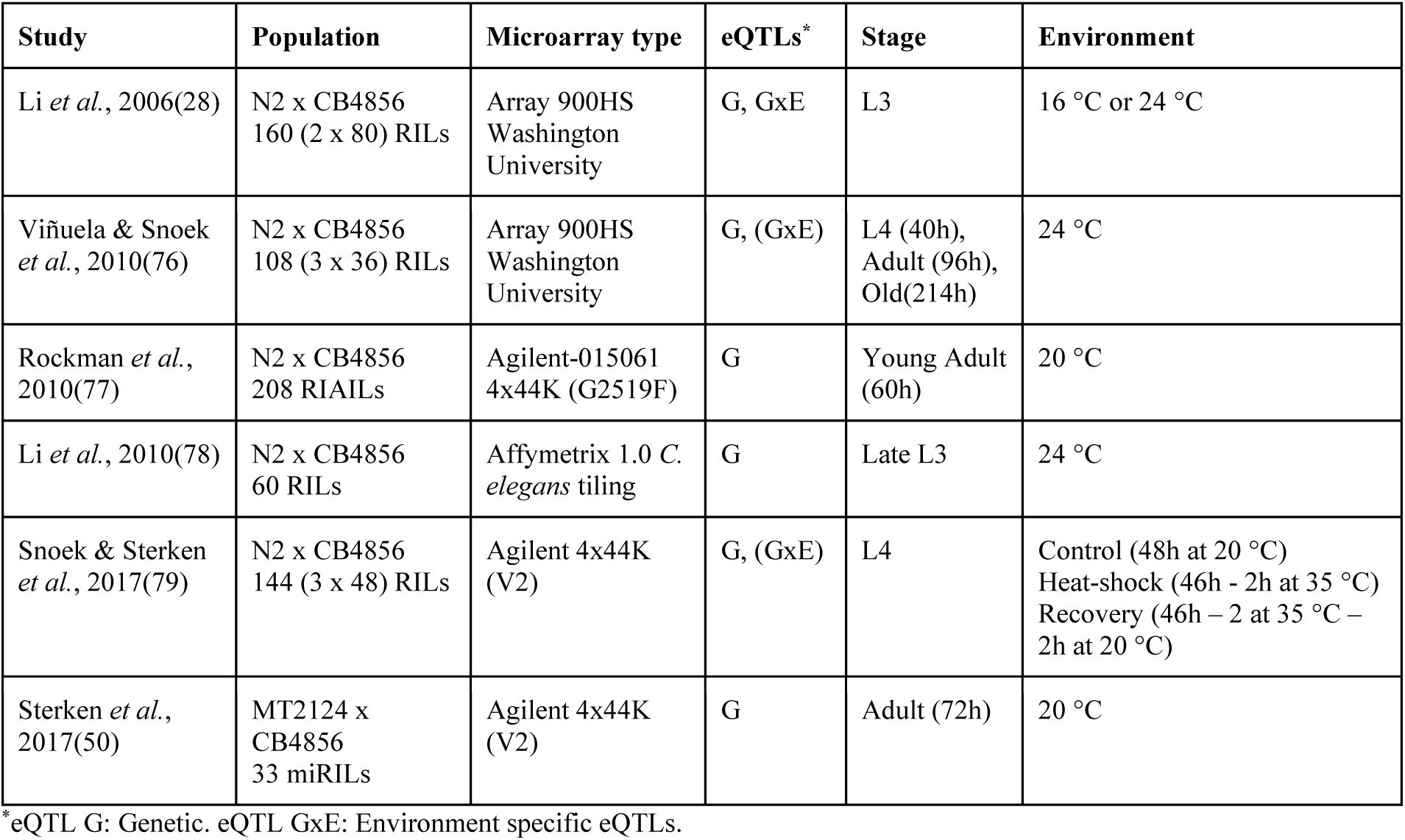
eQTL studies available in WormQTL2.

## Results

### eQTL studies in WormQTL2

WormQTL2 is a browser-based interactive platform and database for investigating expression and other Quantitative Trait Locus (QTL) studies conducted in *C. elegans* (Figure 1). It enables access to the mapping data of six previously published eQTL studies (Table 1) (28,50,76–79). Together, these studies cover over 700 samples, including expression measurements of approximately 20,000 different genes across different life stages and environmental conditions. The effect of genetic variation on gene expression is presented in 11 genome-wide sets of eQTLs from three different RIL populations. The three populations consist of two different CB4856 x N2 populations, recombinant inbred lines (RILs) (28) and recombinant inbred advanced intercross lines (RIAILs) (91, 92) and a mutation introgressed RIL population resulting from a cross between a *let-60* gain-of-function mutant in an N2 background, MT2124, with CB4856 (11, 50). For the Li *et al.* 2006, Viñuela & Snoek *et al.* 2010 and Li *et al.* 2010 studies, the eQTLs were re-mapped with the most recent genetic maps used in Snoek & Sterken *et al.* 2017, which can be obtained from WormQTL2 at the download page accessible by pressing the ‘Download’ button. (27, 79).

**Figure 1:**
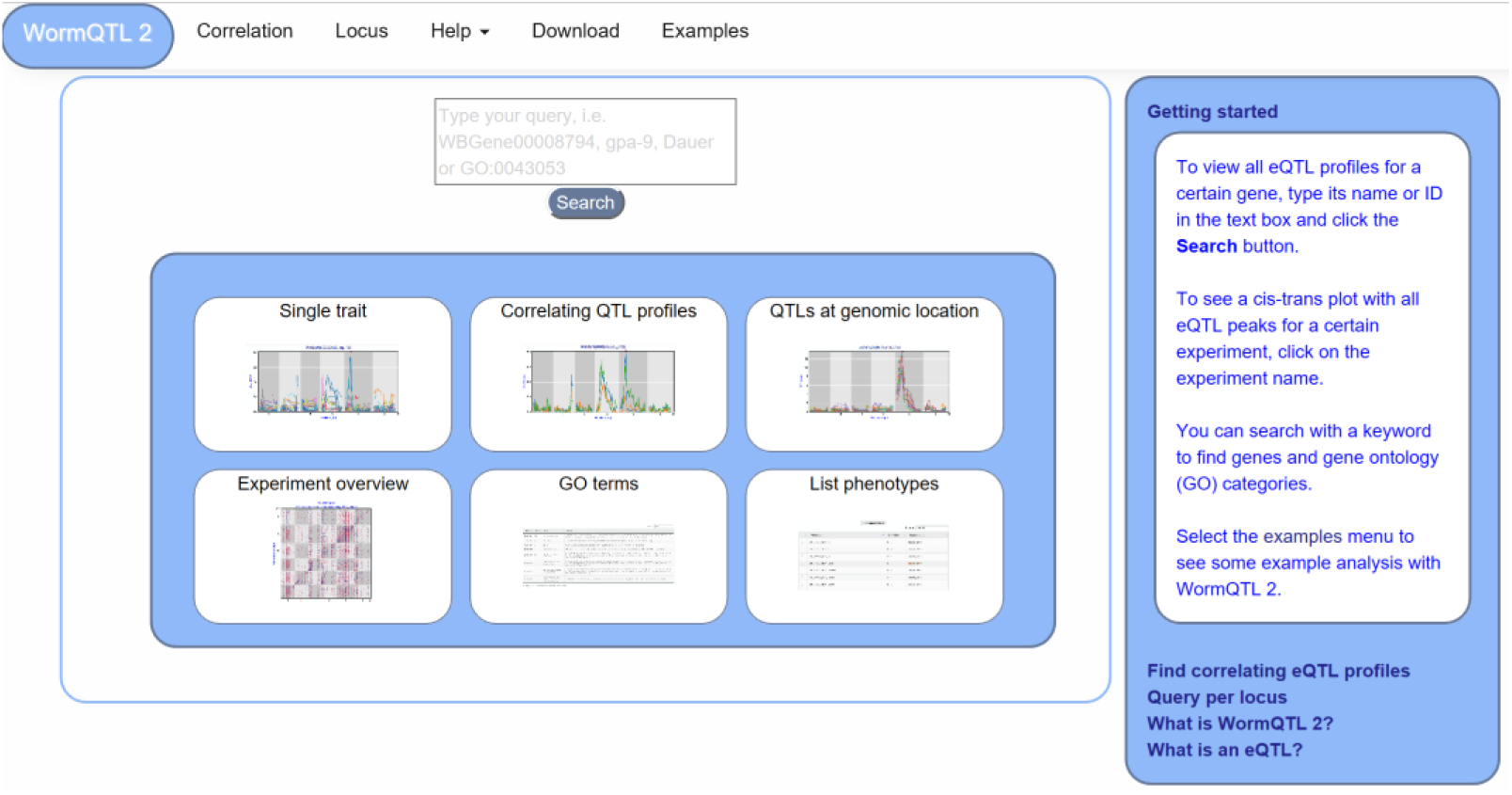
**WormQTL2 Homepage**. On the top of the page the navigation bar can be found. This includes the WormQTL2 logo, which functions as a home button. It also includes a fast link to the Correlation and Locus overviews as well as links for help, data download, and visual examples. The search box is located in the centre, in which genes, phenotypes and GO terms can be entered. Shown in the blue middle square are the buttons for the investigations of single traits, correlating QTL profiles, QTLs at a specific locus, all eQTLs of an experiment, eQTLs per GO term, and a list of all phenotypes for which QTLs can be shown. On the right-side a bar with information about WormQTL2.0 and QTLs in general can be found.

The first eQTL study in *C. elegans* was published in 2006 by Li, *et al.* (28), where variation in gene expression was reported among N2 x CB4856 RILs grown at two different temperatures (16 °C vs 24 °C). In 2010, three eQTL studies in *C. elegans* were published (76–78). Viñuela & Snoek *et al.* showed age specific eQTLs, Li *et al.* investigated variation in splice variants, and Rockman *et al.* used eQTLs to show that phenotypic variation in *C. elegans* is determined by selection on linked sites. These three eQTL studies were used in many follow-up investigations/analyses focusing on how genetic variation affects gene expression, on the regulation of specific genes, or on the molecular pathways underlying phenotypic variation. For instance, a *trans*-band on chromosome X observed by Rockman *et al*. was later identified to result from mild starvation and linked to genetic variation in the *npr-1* gene (3). Snoek & Sterken *et al.* showed the effect of heat-stress and recovery on eQTL distribution and occurrence as well as the contribution of *trans-*eQTLs to cryptic variation (79), and Sterken *et al.* showed the interaction between genetic variation, gene expression and a *let-60* gain-of-function mutation (11, 50). An important overall conclusion drawn from these analyses was that eQTLs are highly dependent on the ambient environment and sensitive to induced background mutations.

Lately, the diversity of molecular phenotypes for which natural variation can be found and used to map QTLs has been expanded to proteins (80, 93) and metabolites (81). The associations of these molecular phenotypes with variation in gene expression, eQTLs and classical phenotypes and QTLs have yet to be explored. For such applications, WormQTL2 provides the data and the interactive platform.

### QTL studies in WormQTL2

WormQTL2 currently provides access to the data and QTLs of 32 RIL-based QTL studies in *C. elegans* (Table 2). These studies include many “classical phenotypes” as well as molecular phenotypes such as metabolite and protein levels. We compiled the list of studies using two reviews listing older studies, mostly pre-2000 (1, 35), and performed a literature search for more recent studies from 2000-onwards. For many studies we could obtain the relevant data from the supplemental information (2,3,5,6,21,34,41,47,49,53,54,57–59,76,80,81,83–86,89–91). Where such data was not provided in journal supplements, approaching the authors was successful for most studies (6-8,11,13,55,82,87,88,91). Data from Bergerac BO x N2 populations was not included as the genetic map only consisted of a few markers and the Bergerac BO strain contained active transposons, complicating the interpretation of the data (35). In summary, at this moment WormQTL2 provides full access to all QTL studies up to 2018, where the required data was available or kindly provided upon request.

The datasets included provide full access to all underlying raw data. For each study a genetic map is available (updated to Wormbase WS258 coordinates), as well as the raw data used for mapping and the output of a single marker model. These data allow users to either access the raw data and run alternative analyses as they wish or access already mapped QTL information. All data were mapped using a single marker model, which is shown in WormQTL2. In total, 929 QTL were mapped for 1091 traits (-log_10_(p) > 3.5). To make these data insightful, trait-names were standardized where possible (e.g. consistent use of ‘body’ in relation to measurements of size, volume, length of the animal body). Furthermore, traits were coupled to gene ontology (GO) terms, to facilitate coupling to transcriptomics data. These curatorial steps greatly facilitate analytical access, increasing the ways in which the user can interact with the data.

### Starting your search using WormQTL2

The homepage offers several approaches to investigate QTL data, including searching for individual traits, correlating QTL patterns and finding traits that have a QTL at a specific locus (Figure 1). Furthermore, six options are provided for quick navigation to specific investigation paths. A detailed description of WormQTL2 navigation and use can be found in the manual (**Supplementary Manual**). In general, the search function can be used to find QTL profiles for one or more traits or genes in one or all experiments. Also, GO-terms can be entered to find all genes annotated with that GO-term and investigate their eQTL profiles. Any search input not directly matching with a gene-ID or GO-term will report the genes and GO-terms with matching descriptions. Divided over several interactively linked pages, different functions are available for investigation and exploration.

### Selecting experiments

Experiments of interest can be selected from the experiment overview page. In this table, basic information about the experiment can be found, such as population used, developmental stage, temperature and source publication. Publications are linked to their PubMed pages for easy access to the experimental details. Data from all experiments, such as QTL profiles, genetic maps and phenotypes are available in WormQTL2 and can be downloaded or directly accessed in flat text format, for instance to further explore with programming languages such as R or Python.

For easy access of the main functionality every WormQTL2 page shows the navigation bar at the top of the page (Figure 1). It can be used for a selection of graphical overviews, investigations and information. To return to the homepage and search function, the “WormQTL2” button in the left upper corner can be used; for each dataset, “correlation” can be used to find traits with correlated QTL profiles; “locus” shows all traits with a QTL at the specified marker or genomic position; and frequently asked questions and other info can be found by clicking “help”. “Examples” leads to an interactive graphical overview of several different functions of WormQTL2.

#### WormQTL2 use cases

WormQTL2 was developed to facilitate meta analyses of QTL and eQTL data for extended investigations and distinguishes itself from other databases by enabling interactive selection of groups of genes or traits based on a common genetic effect. Physical marker positions have been used to integrate the genetic maps of the different populations to enable direct comparisons between eQTLs found in different experiments and populations. Furthermore, WormQTL2 uniquely allows users to find whether a group of genes (such as those sharing a specific GO term) have a shared *trans*-band, a so-called hotspot of eQTLs, which can then be efficiently investigated further for identifying other genes with co-locating eQTLs by the integrative tools. The genes with co-locating eQTLs can then be exported as a list for further investigations or to an external analysis platform. Overall, genes with a shared genetic architecture are easily investigated within and outside WormQTL2, making it a versatile tool for *C. elegans* researchers.

### Example 1: Cryptic variation in gene expression

The environment-specific as well as environment-independent eQTLs for individual or small groups of genes can be easily found by using the search box on the homepage. For instance, gene *gmd-2* has very similar eQTL profiles across experiments, a *cis-*eQTL at chromosome I and a *trans-*eQTL at chromosome V, most prominent in the juvenile and young adult stages (Figure 2) (28,76,77,79). But both *cis*- and *trans-*eQTL are absent from older worms (76, 77) as well as when the genetic background contains a *let-60gf* mutation (50). This shows the hidden/cryptic variation affecting the expression levels of a gene across experiments. This can be very specific, for example *hsp-12.3* has a *cis*-eQTL on chrIV in Rockman *et al.* 2010 and not in any of the other experiments, yet it has a co-locating *trans*-eQTL in both heat stress and recovery conditions in Snoek & Sterken *et al.* 2017.

**Figure 2:**
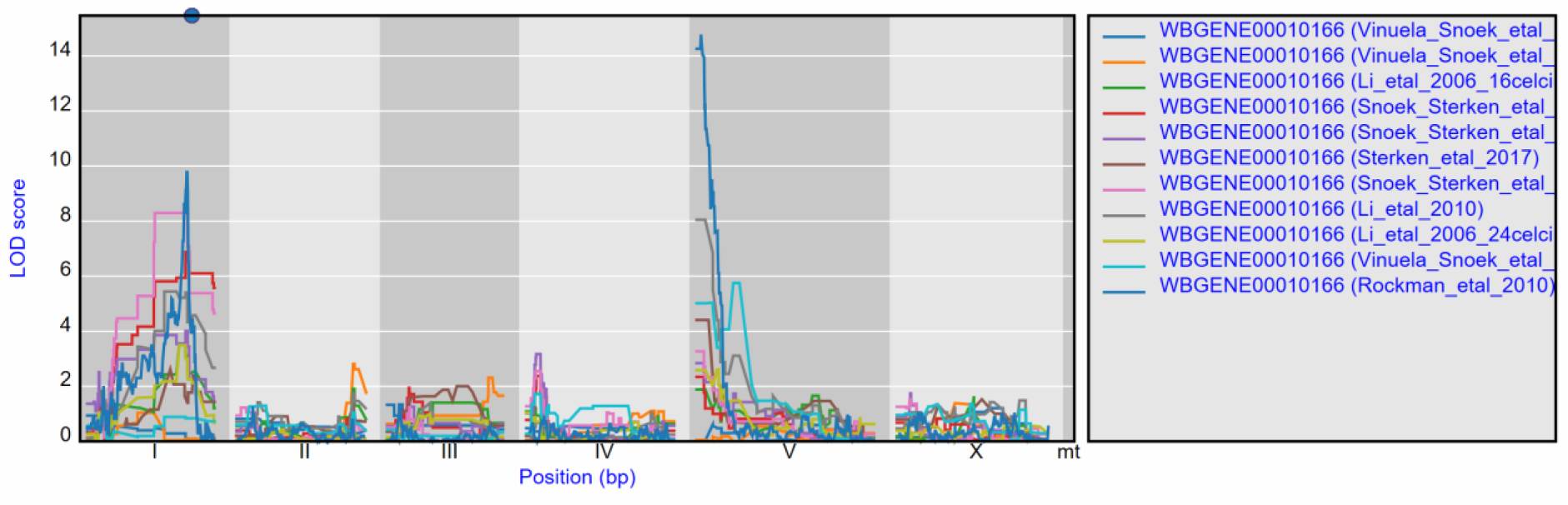
**eQTL profiles of *gmd-2*** from all experiments hosted in WormQTL2. The significance profile (-log_10_(p)) per experiment for *gmd-2* is shown in different colours per experiment. The legend is shown on the right. Chromosomes are indicated by different grey backgrounds and below the x-axis. The web-based plot is interactive; a mouse over provides the exact base pair position of each QTL or other point on the profile.

Similar cryptic variation can also be investigated for other genes, such as *pgp-7*, which has a very prominent *cis-*eQTL on chrX when a *let-60gf* mutation is present in the genetic background (50). Yet in many other experiments it has a *trans*-eQTL on chrV (76,77,79). This shows that different polymorphic regulators exist whose action depends on the developmental stage as well as on the genetic background. The most significant *trans*-eQTL was found in the control conditions of Snoek & Sterken *et al.* 2017 (Table 1); by using WormQTL2’s correlation function on this experiment we can find two *pgp-7* homologs, *pgp-5* and *pgp-6* which also have a *trans*-eQTL at this locus and likely share a common polymorphic regulator (Figure 3).

**Figure 3:**
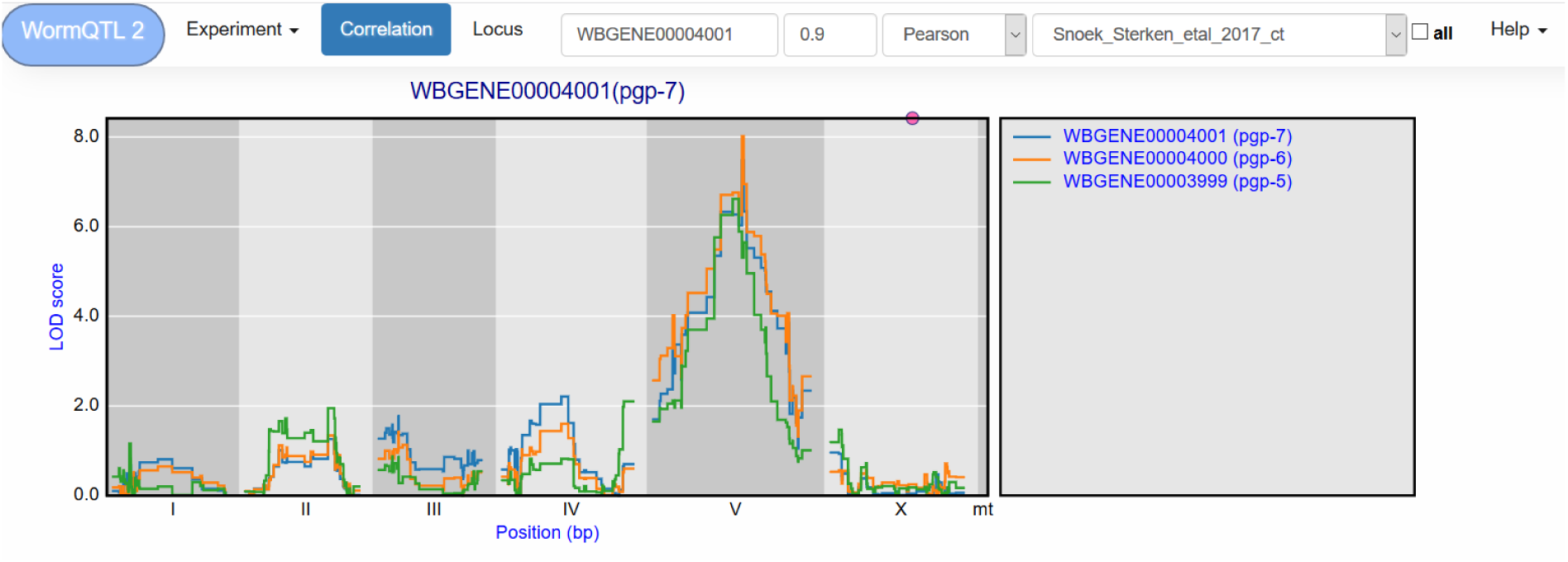
**Correlation between eQTL profiles** of *pgp-7*, *pgp-6* and *pgp-5*. Correlation between the eQTL profile of *pgp-7* and the other genes was set to report those profiles with a Pearson correlation > 0.9, see navigation bar at the top. Genes are shown in different colours. The legend is shown on the right. Chromosomes are indicated by different grey backgrounds and below the x-axis. The web-based plot is interactive; a mouse over provides the exact base pair position of each QTL or other point on the profile.

The correlation between eQTL profiles can show co-regulated groups of genes, the experiment in which they are co-regulated and the regulatory loci involved. When we inspect *daf-18* and the genes with a correlated eQTL profile, we find that *daf-18* is part of a group of co-regulated genes with two regulatory loci only during heat-stress conditions (79), one on chrI and one on chrV (Supplementary Figure 1). Moreover, when the group of genes is enriched for one or more GO terms, a table is provided below the gene table. The *daf-18* co-regulated genes are enriched for larval development, suggesting an effect of heat stress on larval development through *daf-18* expression variation and possibly the loci on chrI and chrV. Comparing the eQTL profiles of genes and groups of genes in different experiments shows the dynamic nature of polymorphic regulatory loci and the genetic architecture underlying cryptic variation.

### Example 2: GO term investigation

Groups of genes can also be selected based on GO terms. Per experiment, the eQTL profiles of all genes annotated with a specific GO term can be shown, with the most significant 15 pre-selected. When, for example, “cell cycle” is entered in the search box, a list of genes and GO-terms is returned. From this list we can pick GO term “regulation of cell cycle” and study the eQTL profiles of the genes involved in this process (Figure 4). In the Viñuela & Snoek *et al.* 2010 juvenile set, 39 genes with co-locating eQTLs can be found on chrI, indicating a polymorphic regulator for the cell cycle can be found at this locus. When we observed the eQTL profiles in other studies the co-location at the locus is gone, indicating the regulator is specific to the juvenile life stage.

**Figure 4:**
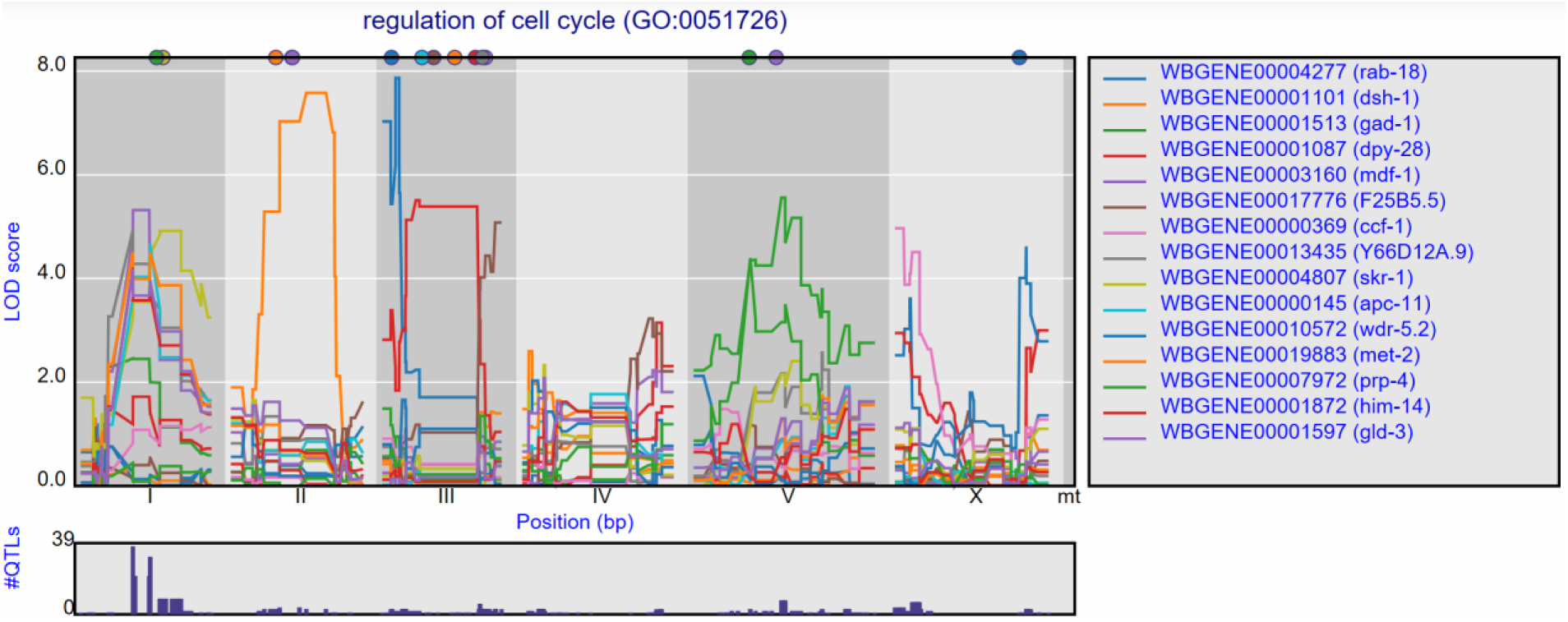
**Co-locating eQTL profiles of genes annotated with GO-term “regulation of cell cycle”**. The eQTL profiles of the top-15 highest eQTLs are shown as profiles. A histogram of all peaks -log_10_(p) > 3.5 is shown below the profile plot to detect possible regulatory hotspots, as can be observed here on chromosome I. Genes are shown in different colours. The legend is shown on the right. Chromosomes are indicated by different grey backgrounds and below the x-axis. The web-based plot is interactive; a mouse over provides the exact base pair position of each QTL or other point on the profile.

This can also be observed for the genes annotated with a related GO term “chromosome segregation” where 41 eQTLs can be found co-locating on chromosome I in the juvenile stage, but not in the reproductive or old stage. In the old stage 14 co-locating eQTLs can be found at chromosome V (Supplementary Figure 2).

These examples show that starting with groups of genes sharing a GO term can be a great start for exploring eQTL data in order to find co-locating eQTLs of genes with a shared function and possibly identify the position of GO term specific polymorphic regulators.

### Example 3: Exploring a trans-band

A *trans*-band can be selected by clicking the histogram under the *cis*/*trans* plot. This leads to a list of genes that pass the significance threshold at the selected marker. For example, in the Rockman *et al.*, 2010 (77) data, marker rmm1258 (Chr X at 3.8 Mb) can be selected, leading to a list of 126 genes at a –log_10_(p) threshold of 3.5. However, this list contains both *cis*- and *trans*-eQTLs. To select the *trans*-eQTLs a minimal distance threshold can be specified to remove genes that are located close to the selected locus. For example, setting this threshold to 2 million base pairs narrows-down the list to 111 genes (Figure 5). These can be investigated further, both within and outside of WormQTL2, to predict their regulator pathway or biological function.

**Figure 5:**
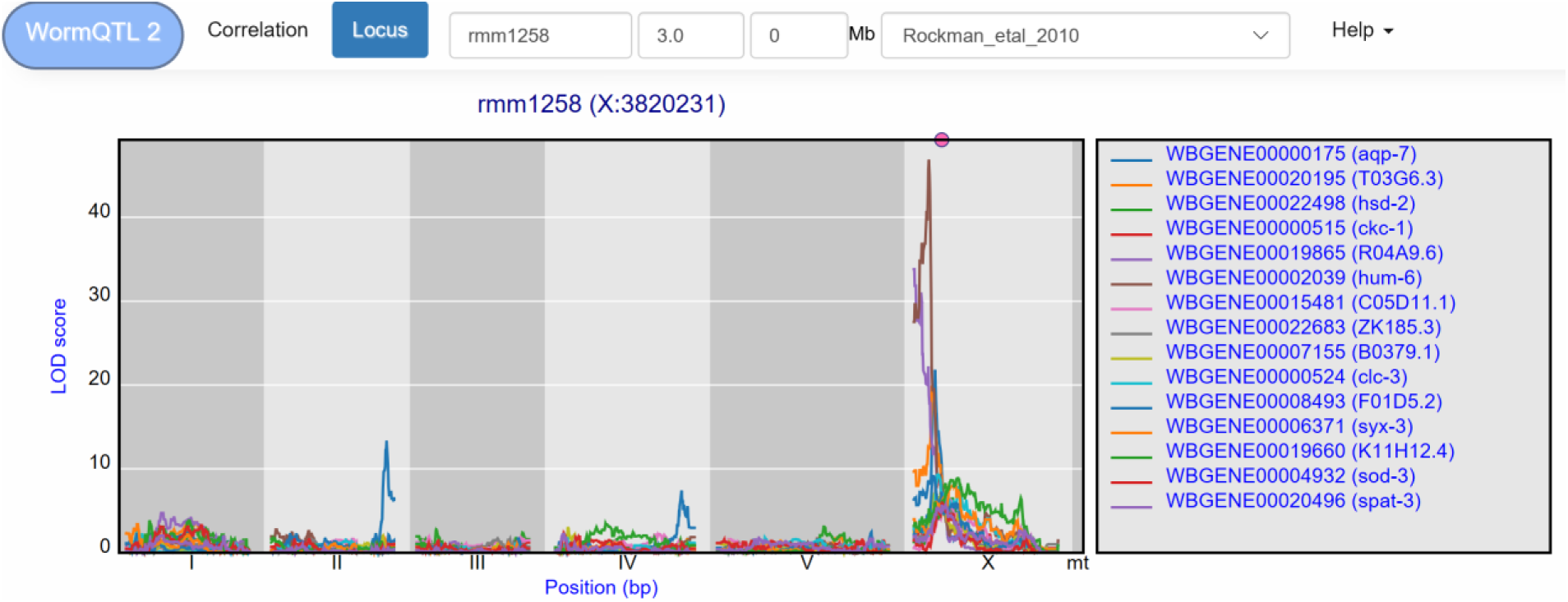
**eQTL transband on chromosome X in Rockman *et al.* 2010.** The eQTL profiles of the top-15 highest eQTLs at marker rmm1258 in Rockman *et al.* are shown as profiles. Colours indicate different genes. The legend is shown on the right. Chromosomes are indicated by different grey backgrounds and below the x-axis. The web-based plot is interactive; a mouse over provides the exact base pair position of each QTL or other point on the profile. Moreover, more than the top-15 can be added to the plot creating a custom overview.

WormQTL2 offers users the option to investigate the eQTL patterns of different studies at the locus location. For instance, it can be informative to determine if a *trans*-band occurs in other studies, by selecting a study from the drop-down menu. WormQTL2 will select and display the eQTLs at the nearest marker for any study selected. When applied to the rmm1258 *trans*-band, by selecting the heat-shock condition of the Snoek & Sterken *et al.*, 2017 (79) study, we find no clear *trans*-band at the corresponding location (PredX3820001), only 6 genes with a *trans-*eQTL. However, in the control condition of this study 26 genes are found to have an eQTL at this position. This is a clear demonstration that this *trans*-band can be environment specific and disappears under heat-shock in L4 stage nematodes.

### Beyond WormQTL2

Using the data stored, explored and selected through WormQTL2, there are several options for further analysis. Using the ‘Download selected trait IDs’-button, a list of wormbase-ID’s can easily be selected and copied. These IDs can subsequently be used in other online *C. elegance* resources, like the Serial Pattern of Expression Levels Locator (SPELL, (94)). For example, the 111 wormbase-IDs from the *trans*-eQTLs can be inserted as query to find gene expression datasets in which the queried genes display co-expression. Visual inspection of the hits identifies three studies with treatment-related variation in this set of genes. One is the original study (77), one is a study on the innate immunity of *C. elegans* and *Pristionchus pacificus* (95), and, most interestingly, a study on an *aptf-1* mutant (96). This mutant shows lower expression of *flp-11*, suggesting that the *trans*-band is somehow related to neuronal activity. From literature, we know this is actually the case, as the underlying gene is *npr-1*, for which a variant (215V in N2) has a neomorphic gain-of-function mutation, making it responsive to both FLP-21 and FLP-18 (reviewed by (1)). There are many other databases that can be consulted similarly for further investigation, such as WormNet (97), WormBase (98), MODENCODE (99)/MODERN (100), Genemania (101), and StringDB (102).

## Discussion

### The WormQTL2 platform for data access

WormQTL2 offers a comprehensive and interactive QTL-data platform for *C. elegans*. It complements and extends existing data analysis and presentation platforms for QTL studies in *C. elegans* like WormQTL (www.WormQTL.org, (74, 75)) and WormQTL-HD (www.WormQTL-HD.org, (73)) and is developed to support *C. elegans* investigators in analyses of natural genetic variation. All data in WormQTL2 are cross-linked which allows for comparative investigation of QTL patterns across studies and phenotypes in a user-friendly interactive way. This can be of great aid as often, published (e)QTL studies do not include the complete QTL profiles. Several years ago, we developed the WormQTL platform to serve as a central repository for these QTL profiles, including the data needed for re-mapping. However, WormQTL lacks interactive tools suitable for further analyses of eQTL profiles, limiting its practical use for direct exploration. In WormQTL2, QTL profiles for genes, metabolites and phenotypes can be viewed and studied interactively. This facilitates integration of different datasets and allows for comparisons which would otherwise be cumbersome and laborious.

WormQTL2 offers access to RIL-based (e)QTL-data in *C. elegans*. Currently, WormQTL2 offers access to all published eQTL datasets and a majority of the published phenotypic QTL datasets. All datasets were curated and the eQTL studies and genetic maps were updated to a recent *C. elegans* genome version (currently the database runs based on WS258). The QTLs of phenotypes of the included studies were re-mapped using a single marker model for uniformity. This does lead to some differences compared to the original study if specific models and mapping procedures were used. However, next to the interactive front-end, the platform also offers access to the raw data, allowing more experienced users to download the data and run custom investigations, extending the reusability of the observations.

The WormQTL2 platform currently limits itself to RIL-based (e)QTL data. Future development will first focus on integrating data from introgression-line (IL) based studies. There is a rich body of published studies utilizing introgression lines for QTL validation, but also as alternative to RIL-based genome-wide studies. Especially the genome-wide N2 x CB4856 IL panel, and a set of chromosome-substitution lines have been used (30, 103). Currently, the *C. elegans* field increasingly makes use of Genome-Wide Association Studies (GWAS) and wild isolates (32,33,40,91). WormQTL2 is currently developing links to genetic variants in QTL regions through the *C. elegans* Natural Diversity Resource (CeNDR) (33).

### Data exploration and analysis through WormQTL2

WormQTL2 offers the user the possibility to compare (e)QTL patterns across studies. For example, *cis*-eQTLs are associated with polymorphisms in or near the gene itself. Furthermore, *trans*-eQTLs can be represented by so-called ‘*trans*-bands’ or ‘eQTL hotspots’, where a polymorphic regulatory gene (or genes) explains variation in transcript abundance of many genes. From literature across species, it is currently clear that *cis*-eQTLs (i) can result from hybridization differences when using microarrays (77,104–106), (ii) explain more variation than single *trans*-eQTL (77, 107), (iii) are constitutively found across experiments using the same populations and environments (28,71,76,78,79,108), and (iv) are often found for polymorphic genes (28,65,71,77). In contrast, *trans*-eQTLs are strongly environment dependent and seem in large part unique across environments (28,71,76,109).

Comparative analysis of eQTL profiles does come with some (inherent) limitations. The main limitation is platform-based. WormQTL2 offers access to eQTL studies from four different microarray platforms. As microarray technology uses probes on a glass-slide, false negatives may occur due to absence of a probe, preventing the interrogation for that gene (77, 104). Closely related genes can cross-hybridize due to probe similarity (although we try to minimize the risk by excluding probes with multiple blast-hits). Furthermore, false positive eQTLs could be obtained when hybridisation differences due to sequence polymorphisms are mistaken for transcript abundance variation. These QTLs however, can be used as genetic markers or to detect wrongly labelled samples (105,106,110). Hence, users should be mindful about technical limitations when comparing results from different experiments. Nevertheless, for most eQTLs and general patterns cross-platform comparisons can be insightful and useful (50).

Comparison of (e)QTL studies through WormQTL2 depends on the mapping populations involved. We offer analytical access to studies widely differing in statistical power and RIL population. For example, the number of RILs used per (e)QTL dataset ranges from 36 for each of the three conditions in Viñuela & Snoek *et al.*(76), up to 200 in Rockman *et al.*(77) and Snoek *et al.*(34) or even more in the CeMEE panel of Noble *et al.* (111). Furthermore, the size of the genetic map (in centimorgans) of the N2 x CB4856 RIAIL populations is larger than the N2 x CB4856 RIL population (27,28,77). Genetic maps of the mutation included RIL populations and the N2 x DR1350 populations include areas without genetic variation (or information on the genotype) (11,50,88), whereas a multi parental RIL panel contains multiple SNP distribution patterns (34). Finally, the number of markers used to genotype RILs are different between mapping populations. In WormQTL2 we therefore present the eQTL profiles of re-mapped studies, so the best comparisons can be made.

#### Future developments

WormQTL2 aims to provide re-mapped (e)QTLs in *C. elegans*. Currently, re-mapping has been done using a single marker model, making output comparable across studies. As all the relevant data for mapping are hosted, it is possible to integrate alternative models or integrate analysis of different experiments in one (e)QTL mapping model. Genetic maps can also be improved by including gene expression markers (50,79,105,106,110), through sequencing (27) or use of RNA-seq (34, 112). This will lead to eQTLs with a higher resolution and better regulatory prediction. Easy access to the data already enables an efficient start for further exploiting eQTLs and other system genetics data by anyone. In future updates of WormQTL2 these QTL mapping functions can be implemented.

Combining established high-throughput measurement techniques like next generation sequencing (27,32–34), proteomics (80, 93), metabolomics (81) and phenomics (82,83,113) offers great potential for further quantitative genetic analyses across different levels. This wealth of data makes the storage, access and especially the generation of useful and meaningful connections within and between the different types of data increasingly important. Moreover, results from different types of mapping populations can be included. In this way the advantages of IL populations (30,54–56,103,114,115), RIL populations (28,50,92), multi-parental mapping panels (34, 111) and sets of wild-isolates for GWAS (33) can be combined. With this in mind, the next steps for WormQTL2 will be linking eQTL data to polymorphisms from massive sequencing projects of many different ecotypes (32, 33) and including eQTLs and SNPs obtained from RNA-seq experiments. When stored in, and visualised from, the same platform, the SNPs and phenotypes enable the integration of QTL mapping and GWAS investigation, further increasing the detection power of both methods. For example, eQTL datasets have been successfully combined with results from transcriptomic GWAS (116) and allele-specific expression RNA-seq experiments (109).

New tools for investigation and visualisation will be developed in a modular fashion for easy integration and deployment within WormQTL2. Annotations can be expanded beyond GO terms, for example with pathway knowledge, *e.g.* as available through the Kyoto Encyclopaedia of Genes and Genomes (KEGG; www.genome.jp/kegg/), or with gene association networks like WormNet (97), StringDB (102) and Genemania (101). To further investigate the relation between genotype and phenotype variation, WormQTL2 will be expanded with published and new classical/phenotypic QTLs. This enables searching for the possible molecular components underlying a QTL for a specific phenotype and finding the causal genes. Combining highly detailed molecular data, such as generated and shared by the modENCODE consortium (99,117,118), like transcription factor- and histone-binding sites or protein-protein interactions will allow for even more powerful analyses.

WormQTL2 has been designed to easily store and share upcoming RNA-seq data and eQTLs from this data, QTLs from metabolomics and proteomics and visualise and analyse these together. Comparing sets of genes through functional enrichment will enable an even better, more targeted, approach in candidate gene selection and network generation to link gene expression, genetic variation and function. In the near future, tools will be developed to investigate genetic variation in a more systematic, genome- and population-wide manner, enabling more complete and higher resolution system genetics.

We believe the (e)QTL data in WormQTL2 will greatly benefit the *C. elegans* research community, providing a rich source of genetic interactions specifically to worm biologists and geneticists in general. WormQTL2 will serve as a solid platform for in-depth analysis of these interactions to help chart the *C. elegans* gene regulatory networks.

## Experimental procedures

### Transcriptome data

Datasets of the six eQTL experiments were retrieved from GEO or ArrayExpress: GSE5395 (28), GSE15778(78), GSE17071 (76), GSE23857 (77), E-MTAB-5779 (79), E-MTAB-5856 (50). The platform data were also obtained from GEO or ArrayExpress: GPL4043 (28, 76), GPL5634 (78), GPL7727 (77), A-MEXP-2316 (50, 79). The microarray probes were re-mapped against WS258 using blastn (version 2.6.0, win x64(119)). Probes with multiple high-ranking matches to different genes were censored.

### Phenotypic data

The publications reporting *C. elegans* phenotypic QTLs were used to acquire phenotype and genotype data required for QTL mapping. This was done by taking the data directly from separated supplementary information or by contacting the authors. We curated data from 32 publications, comprising 1091 traits (Table 2). For each publication the raw trait data per strain, the genetic map and the QTL data are made available.

Using the obtained phenotypes and the most updated genotype data, the QTLs for each study were re-mapped using a linear single marker model

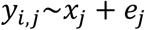

where *y* is the phenotype value of RIL *j* based on the function of marker genotype *i*. Subsequently, a -log_10_(*p*-value) of 3.5 from each marker analysis was used as the threshold to determine the significant QTL shown in table 2. This analysis was performed using a custom-made script in “R” (version 3.4.4, win x64).

**Table 2:** QTL studies available in WormQTL2.

| Paper reference | Title | Parental lines | Type | Traits | QTLs<br>$-\log_{10}(p) > 3.5$ |
| --- | --- | --- | --- | --- | --- |
| Andersen *et al.*, 2014 (3) | A variant in the neuropeptide receptor npr-1 is a major determinant of *Caenorhabditis elegans* growth and physiology. | N2 x CB4856 | RIAIL | 7 | 10 |
| Andersen *et al.*, 2015 (82) | A Powerful New Quantitative Genetics Platform, Combining *Caenorhabditis elegans* High-Throughput Fitness Assays with a Large Collection of Recombinant Strains. | N2 x CB4856 | RIAIL<sup>1</sup> | 34 | 27 |
| Balla *et al.*, 2015 (41) | A wild *C. elegans* strain has enhanced epithelial immunity to a natural microsporidian parasite | N2 x CB4856 | RIAIL | 1 | 0 |
| Bendesky *et al.*, 2011 (8) | Catecholamine receptor polymorphisms affect decision-making in *C. elegans*. | N2 x CB4856 | RIAIL | 1 | 2 |
| Bendesky *et al.*, 2012 (7) | Long-range regulatory polymorphisms affecting a GABA receptor constitute a quantitative trait locus (QTL) for social behavior in *Caenorhabditis elegans*. | N2 x CB4856 | RIAIL | 2 | 1 |
| Evans *et al.*, 2018 (83) | Shared Genomic Regions Underlie Natural Variation in Diverse Toxin Responses. | N2 x CB4856 | RIAIL<sup>1</sup> | 384 | 590 |
| Duveau & Felix, 2012 (47) | Role of pleiotropy in the evolution of a cryptic developmental variation in *Caenorhabditis elegans*. | N2 x AB1 | RIL | 3 | 3 |
| Elvin & Snoek *et al.*, 2011 (49) | A fitness assay for comparing RNAi effects across multiple *C. elegans* genotypes. | N2 x CB4856 | RIL | 108 | 20 |
| Gaertner *et al.*, 2012 (84) | More than the sum of its parts: a complex epistatic network underlies natural variation in thermal preference behaviour in *Caenorhabditis elegans*. | N2 x CB4856 | RIAIL | 5 | 5 |
| Gao & Sterken *et al.*, 2018 (81) | Natural genetic variation in *C. elegans* identified genomic loci controlling metabolite levels. | N2 x CB4856 | RIL | 378 | 154 |
| Glater *et al.*, 2014 (85) | Multigenic natural variation underlies *Caenorhabditis elegans* olfactory preference for the bacterial pathogen *Serratia marcescens*. | N2 x CB4856 | RIAIL | 2 | 2 |
| Greene *et al.*, 2016 (86) | Regulatory changes in two chemoreceptor genes contribute to a *Caenorhabditis elegans* QTL for foraging behaviour. | MY14 x CX12311 | RIL<sup>2</sup> | 2 | 1 |
| Gutteling *et al.*, 2007 (58) | Environmental influence on the genetic correlations between life-history traits in *Caenorhabditis elegans*. | N2 x CB4856 | RIL | 6 | 2 |
| Gutteling *et al.*, 2007 (59) | Mapping phenotypic plasticity and genotype-environment interactions affecting life-history traits in *Caenorhabditis elegans*. | N2 x CB4856 | RIL | 18 | 5 |
| Harvey *et al.*, 2008 (88) | Quantitative genetic analysis of life-history traits of *Caenorhabditis elegans* in stressful environments. | N2 x DR1350 | RIL | 26 | 4 |
| Harvey, 2009 (87) | Non-dauer larval dispersal in *Caenorhabditis elegans*. | N2 x DR1350 | RIL | 2 | 2 |
| Kammenga *et al.*, 2007 (5) | A *Caenorhabditis elegans* wild type defies the temperature-size rule owing to a single nucleotide polymorphism in *tra-3*. | N2 x CB4856 | RIL | 1 | 1 |
| Large, *et al.*, 2016 (6) | Selection on a Subunit of the NURF Chromatin Remodeler Modifies Life History Traits in a Domesticated Strain of *Caenorhabditis elegans*. | LSJ2 x CX12311 | RIL | 5 | 7 |
| Lee *et al.*, 2017 (89) | The genetic basis of natural variation in a phoretic behaviour. | N2 x CB4856 | RIAIL<sup>1</sup> | 1 | 1 |
| McGrath *et al.*, 2009 (13) | Quantitative mapping of a digenic behavioural trait implicates globin variation in *C. elegans* sensory behaviours. | N2 x CB4856 | RIAIL | 2 | 4 |
| Nakad *et al.*, 2016 (53) | Contrasting invertebrate immune defence behaviours caused by a single gene, the *Caenorhabditis elegans* neuropeptide receptor gene *npr-1*. | N2 x CB4856 | RIL | 24 | 17 |
| Noble *et al.* 2015 (2) | Natural Variation in *plep-1* Causes Male-Male Copulatory Behavior in *C. elegans*. | QG5 x QX1199 | RIL<sup>3</sup> | 5 | 5 |
| Rockman & Kruglyak, 2009 (91) | Recombinational Landscape and Population Genomics of *Caenorhabditis elegans*. | N2 x CB4856 | RIAIL | 7 | 6 |
| Rodriguez *et al.*, 2012 (57) | Genetic variation for stress-response hormesis in *C. elegans* lifespan. | N2 x CB4856 | RIL | 9 | 1 |
| Schmid *et al.*, 2015 (11) | Systemic Regulation of RAS/MAPK Signaling by the Serotonin Metabolite 5-HIAA | MT2124 x CB4856 | RIL<sup>4</sup> | 2 | 3 |
| Singh *et al.*, 2016 (80) | Natural Genetic Variation Influences Protein Abundances in *C. elegans* Developmental Signalling Pathways. | N2 x CB4856 | RIL | 10 | 1 |
| Snoek *et al.*, 2019 (34) | A multi-parent recombinant inbred line population of *C. elegans* allows identification of novel QTLs for complex life history traits. | JU1511 x JU1926 x JU1931 x JU1941 | mpRIL<sup>5</sup> | 21 | 72 |
| Snoek & Orbidans *et al.*, 2014 (55) | Widespread genomic incompatibilities in *Caenorhabditis elegans*. | N2 x CB4856 | RIL | 4 | 3 |
| Stastna *et al.*, 2015 (54) | Genotype-dependent lifespan effects in peptone deprived *Caenorhabditis elegans*. | N2 x CB4856 | RIL | 2 | 2 |
| Viñuela & Snoek *et al.*, 2010 (76) | Genome-wide gene expression regulation as a function of genotype and age in *C. elegans*. | N2 x CB4856 | RIL | 2 | 0 |
| Zdraljevic *et al.*, 2017 (23) | Natural variation in a single amino acid substitution underlies physiological responses to topoisomerase II poisons. | N2 x CB4856 | RIAIL<sup>1</sup> | 2 | 3 |
| Zhu *et al.*, 2015 (90) | Compatibility between mitochondrial and nuclear genomes correlates with the quantitative trait of lifespan in *Caenorhabditis elegans*. | N2 x CB4856 | RIAIL | 15 | 11 |

<sup>1</sup> Supplemented by an QX1430 x CB4856 cross; <sup>2</sup> CX12311 is N2 without *npr-1* and *glb-5*; <sup>3</sup> QG5 is *him-5* (e1490) > AB2,
QX1199 is *him-5* (e1490) > CB4856; <sup>4</sup> MT2124 is a *let-60* gain-of-function mutant; <sup>5</sup> multi (four) parental cross.

### Genetic maps

For each population the most detailed genetic map available was used for remapping the eQTL. For the experiments on the N2 x CB4856 RILs this was a low-coverage sequencing based genetic map (27) consisting of 729 markers. As not all RILs in this population were sequenced, the genotypes of those RILs were imputed (79). For the N2 x CB4856 RIAIL population a SNP-based genetic map consisting of 1454 markers was used (91). The genetic map of the MT2124 x CB4856 population consists of an expanded FLP-map with 247 markers (11,27,50). The marker locations of each map were updated to WS258

For the phenotype QTL studies, the genetic map used in re-mapping can also be found on WormQTL2. This includes five additional genetic maps. First, the map for the expanded RIAIL set, which added 359 strains to the panel, without the N2 *npr-1* allele and a transposon insertion to reduce the effect of *peel-1* (82). Second, the JU605 x JU606 RILs, which are made from N2 with a *let-23(sy1)* mutation crossed with an AB1-genetic background introgression line with an N2-segement containing the *let-23* mutation (47). Third, the MY14 x CX12311 RILs, where CX12311 is an N2 strain with the wild-type *npr-1* and *glb-5* alleles (86). Fourth, RILs from an N2 x DR1350 cross (88). Fifth, RILs from a QG5 x QX1199 cross, where QG5 is the strain AB2 and QX1199 is CB4856, both carrying a *him-5(e1490)* mutation (2). Also, these maps were updated to WS258 coordinates.

### Microarray normalization and processing

For each array-type recommended normalization methods were used. Each study was normalized independently (120). For all the normalization procedures the limma-package from Bioconductor (121) was used in “R” (version 3.4.2, win x64). The array data of the two studies based on the GPL4043 platform were background corrected using the *subtract* method, the within-array normalization method used was *printtiploess*, and between-array normalization method used was *quantile*. The array data of the tiling-array (GPL5634) (78) was reprocessed from the raw data and was batch corrected to remove between batch effects. Thereafter the tiling array data was summarized per gene using the *quantile* function in “R”. All five quantiles were used as input for subsequent analyses. For the two Agilent platforms (GPL7727 and A-MEXP-2316) no background correction was applied (122), the within-array normalization method used was *loess*, and the between-array normalization method used was *quantile*.

After normalization, the intensities were log_2_-transformed for subsequent analyses. The Li *et al.*, 2006 experiment on the GPL4043 platform was corrected for array-specific effects due to a heterogeneous hybridization environment. In order to remove this effect, the difference between the array and the total average over all arrays was subtracted from the samples. The Viñuela & Snoek *et al.*, 2010 experiment on the GPL4043 platform did not suffer from such an artefact.

### Wrongly labelled samples

In order to reduce the noise in re-mapping, the correlation between the transcriptome profiles and the linkage map of the population used was determined based on known *cis*-eQTL in *C. elegans* (79, 105). If switched labels were detected, these were corrected and if no fitting correlations could be made, samples were censored. The datasets with correct strain labels and normalized values were made available for downloading and were used to produce the genotype split plots.

### eQTL mapping

For each study the eQTL were re-mapped using a single marker model, as in (50, 79). In short, the linear single marker model

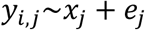

was used, where the log_2_-normalized intensity (*y*) of spot *i* of RIL *j* was explained over the genotype at marker location *x* of RIL *j*. Per gene, the spot with the highest significance was selected as representative. The significance of the correlation and its effect were used to produce the *cis*/*trans* eQTL plots of the study and the detailed eQTL profile per gene.

### Threshold determination

In WormQTL2 the user can set the threshold for determining if an eQTL is significant as this threshold can be dependent on many factors or even user preference. However, the default set thresholds are those used in the original papers. -log_10_(p) thresholds for Li *et al.* 2010: 4.2, Viñuela & Snoek *et al.* 2010: 3.8, Rockman *et al.* 2010: 4.5, Snoek & Sterken *et al.* 2017: control 3.9, heat shock 3.9, recovery 3.9 and Sterken *et al.* 2017: 3.5 are used as the default settings.

### QTL correlation analysis

All pairwise correlations between eQTL patterns were calculated using the Pearson correlation coefficients between the -log_10_(p) values of the eQTL patterns of genes within an experiment using a custom python function (**Supplemental script**). Searching for genes with an eQTL at a specific locus is implemented by selecting genes that have a -log_10_(p) score above the given threshold at the marker closest to the specified locus. To select *trans-*eQTLs, genes with a *cis*-eQTL can be excluded based on their physical distance to this marker.

### Additional data and webpage development

GO terms were downloaded from WormBase and gene descriptions from Ensembl BioMart (123). WormQTL2 was developed using the Python Django web framework. The backend runs on an Ubuntu 17.10 Linux server, using the Apache web server version 2.4.27 and a MySQL 5.7.13 database. The web frontend is implemented via Django templates in HTML and Javascript, using the D3 library and Jquery. The *cis*/*trans* plot and QTL profile plots build upon work by Karl Broman (124).

### Legacy data hosted at WormQTL1

Transcriptomics and genomics data from three papers that was hosted on WormQTL1, that was no hosted anywhere else and that did not fall under the category QTL experiments, was submitted to ArrayExpress. This concerns data from an experiment investigating genetic variation in 48 *C. elegans* isolates (E-MTAB-8126) and gene expression variation in these isolates (E-MTAB-8132) (38), transcriptomics data from an experiment comparing N2 and a *bar-1* loss-of-function mutant (EW15; ga80) (E-MTAB-8128) (125), and transcriptomics data from an experiment growing eight strains on two different bacteria (E-MTAB-8129) (126).

## Acknowledgements

BLS was funded by Netherlands Organisation for Scientific Research (project no. 823.01.001). The authors want to thank the Kammenga lab members for testing and commenting on beta versions of the website. The authors also want to thank Patrick McGrath, Andrés Bendesky, Cori Bargmann, Simon Harvey, Matt Rockman, and Erik Andersen for sharing raw data of their publications.

## Author contributions

HN and BLS came up with the idea for WormQTL2. HN, MGS and BLS designed WormQTL2. HN wrote the code for WormQTL2. MGS, BLS, AJvZ and MH managed the data. DR and JEK provided resources. BLS, MGS, HN wrote the manuscript, with contributions of all co-authors.

## Supplementary Figures

**Supplement figure 1:**
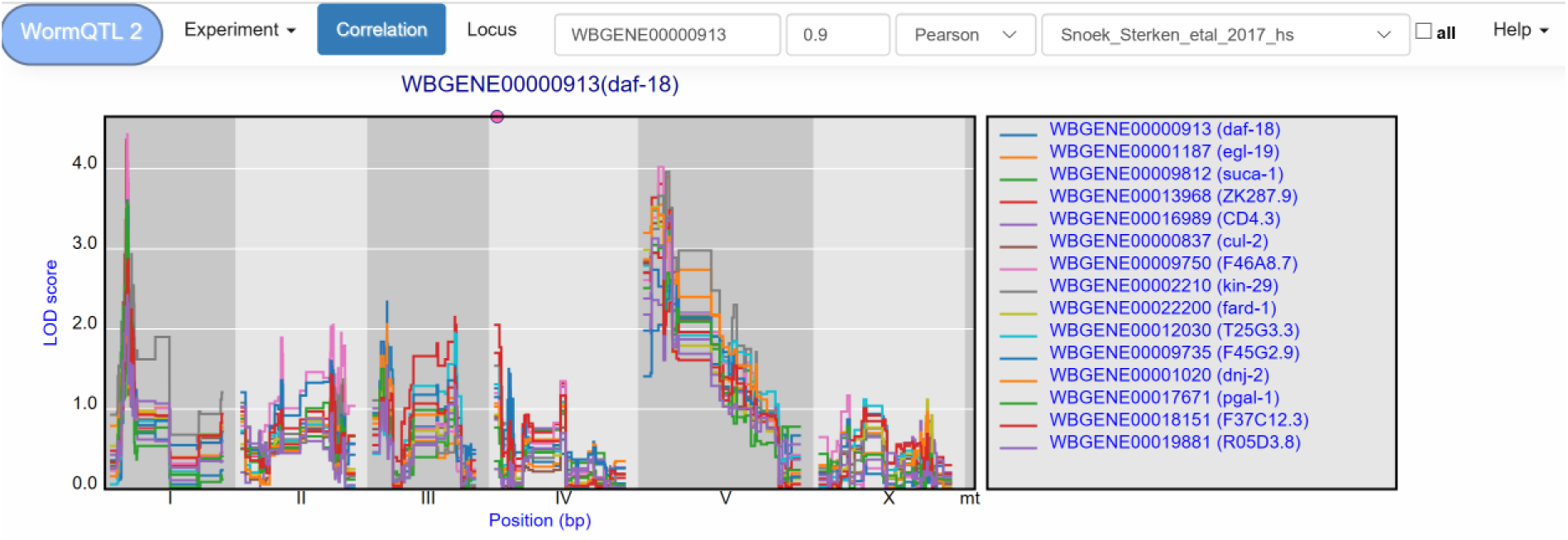
eQTL profiles of genes with a highly correlated eQTL profile to the daf-18 eQTL profile.

**Supplementary Figure 2:**
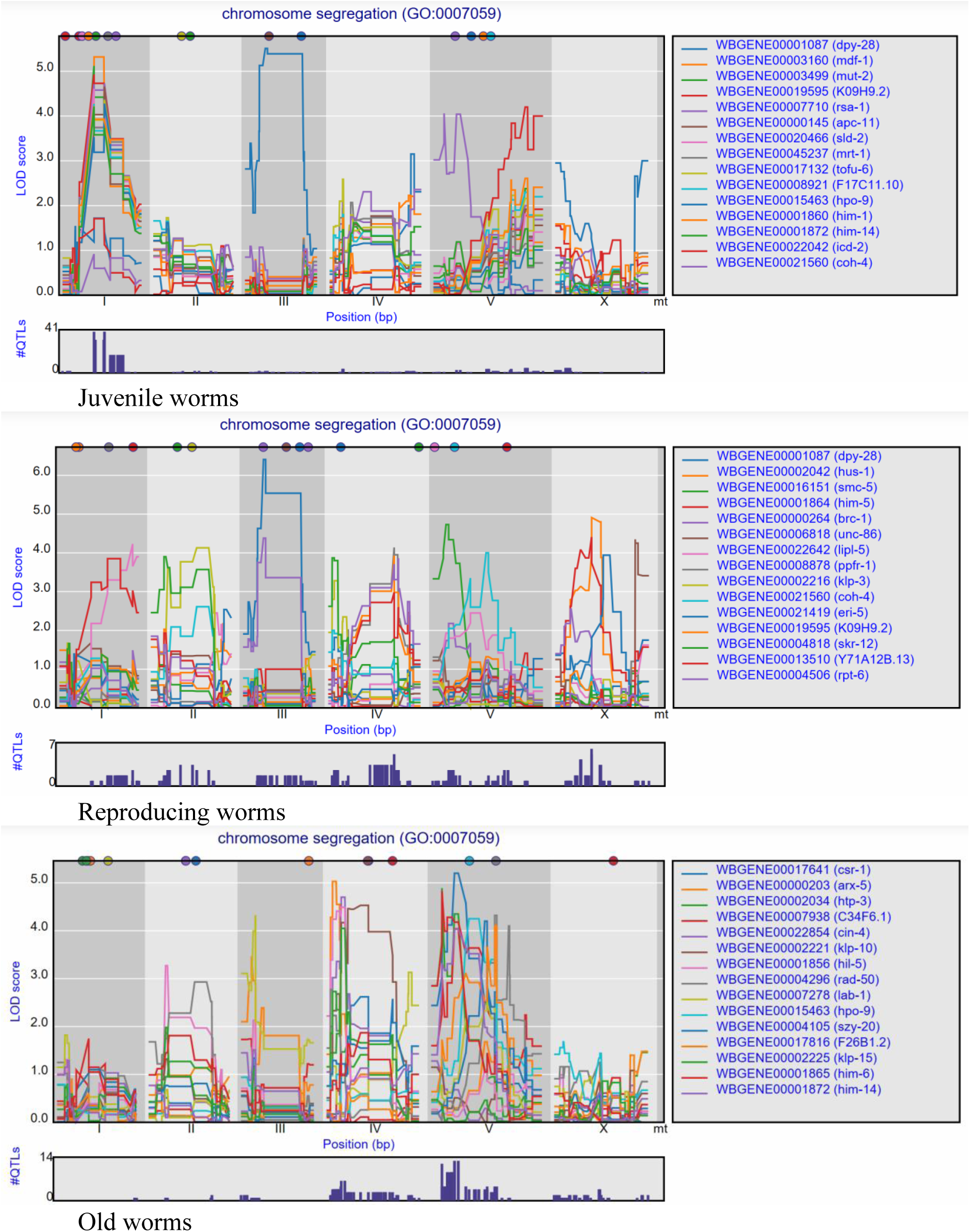
eQTLs for genes with GO-term “chromosome segregation”, in three different life-stages.

**Supplementary Manual** See WormQTL2 website: www.bioinformatics.nl/WormQTL2/

